## Supplementary Information for "Multimodal *in vivo* recording using transparent graphene microelectrodes illuminates spatiotemporal seizure dynamics at the microscale"

### Optimization of graphene microelectrode array fabrication

Our graphene device fabrication follows closely the protocol outlined by Park et al. in 2016, with several important modifications[1]. Firstly, monolayers of graphene grown on copper foils via chemical vapor deposition (CVD) were transferred to the wafer using a bubble transfer method similar to that described by Gao et. al. in 2012[2]. While conventional graphene transfer methods rely on complete etching of metal substrates in suitable etchants, which causes damage to the graphene and leaves metal residues, bubble transfer using electrolysis of water enables transferring large-area graphene films with high purity and without inducing cracking or damage. Initially, we fabricated devices with either one or two stacked monolayers of graphene to assess the relationship between number of graphene layers and electrochemical impedance (Supplementary Fig. S2a). While we found that adding a second layer of graphene reduced the impedance (2.16 ± 0.23 MΩ for 2-layer vs. 2.75 ± 0.26 MΩ for 1-layer graphene at 1 kHz), we sought to reduce the impedance further without adding additional graphene layers, as these would reduce the optical transparency of our devices. To this end, we explored chemical doping methods for enhancing the out-of-plane conductivity of graphene.

Exposing graphene to nitric acid ($HNO_3$) results in the adsorption of electropositive $NO_3^-$ groups on the graphene surface, a p-type or hole doping, which has been shown to reduce electrochemical impedance and improve the noise characteristics of graphene electrodes[3–5]. We optimized a process for $HNO_3$ doping of graphene, and performed this doping layer-by-layer after the transfer of each graphene sheet to the wafer substrate. Layer-by-layer doping has been shown to produce doped graphene sheets with reduced and more stable sheet resistance values compared to doping after stacking of graphene monolayers, where only the topmost layer experiences doping[3, 6]. To assess the performance of devices prepared under different conditions, we compare impedance values both at the commonly-used 1 kHz reference frequency, and at the lower frequency of 100 Hz, since many physiologic signals of interest in the brain are of lower frequency content than the typical 1 kHz reference frequency (Supplementary Fig. S2b,c). We find that the impedance relationships between 1-layer, 2-layer, and doped 2-layer graphene devices are similar at 1 kHz and 100 Hz, with the advantage of chemical doping being more pronounced at lower frequencies: the 100 Hz impedance of 1-layer devices is 25.6 ± 11 MΩ, 2-layer devices is 15.9 ± 5.5 MΩ, and $HNO_3$-doped devices is 2.07 ± 0.65 MΩ. The significant spread of 100 Hz impedance values for the un-doped device preparations can be attributed to their high impedance in this lower frequency range, which limits the accuracy of the EIS measurement. As a low electrode impedance is essential for obtaining recordings with high signal-to-noise ratio (SNR)[7, 8], these results indicate that $HNO_3$ doping is a useful approach for improving the performance of graphene microelectrodes, especially at lower frequencies where most physiologic signals of interest occur. In addition to EIS measurements, cyclic voltammetry (CV) measurements were taken to assess the effect of $HNO_3$ doping on the charge storage capacity of graphene electrodes. Doping with nitric acid increased the charge storage capacity from 22.87 $\mu C/cm^2$ to 64.44 $\mu C/cm^2$, indicating that this method of doping may also improve the performance of graphene electrodes for neural stimulation applications (Supplementary Fig. S2d).

**Supplementary Figures:**

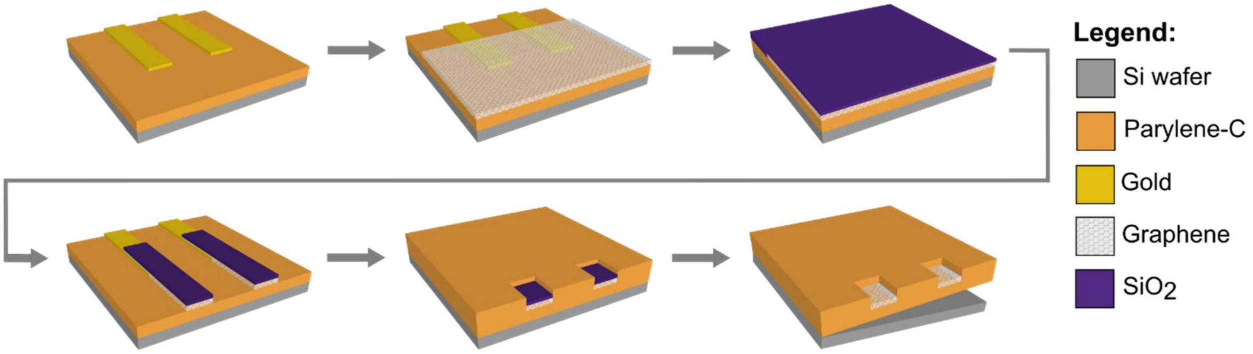


**Supplementary Fig. S1 | Fabrication schematic for transparent graphene micro-ECoG.** Details of the fabrication are given in the Methods section. Briefly, parylene-C is deposited onto a Si wafer and Ti/Au traces are patterned on the parylene-C. CVD-grown graphene is transferred to this substrate using a bubble transfer method. An $SiO_2$ layer is deposited to protect the graphene during later processing steps, then the graphene and $SiO_2$ layers are patterned and etched. A top layer of parylene-C is deposited to encapsulate the device, then VIAs are etched to open the graphene electrode contacts as well as open the Ti/Au bonding pads at the back end of the device. Finally, the $SiO_2$ covering the graphene electrode contacts is chemically etched away and the devices are removed from the wafer.

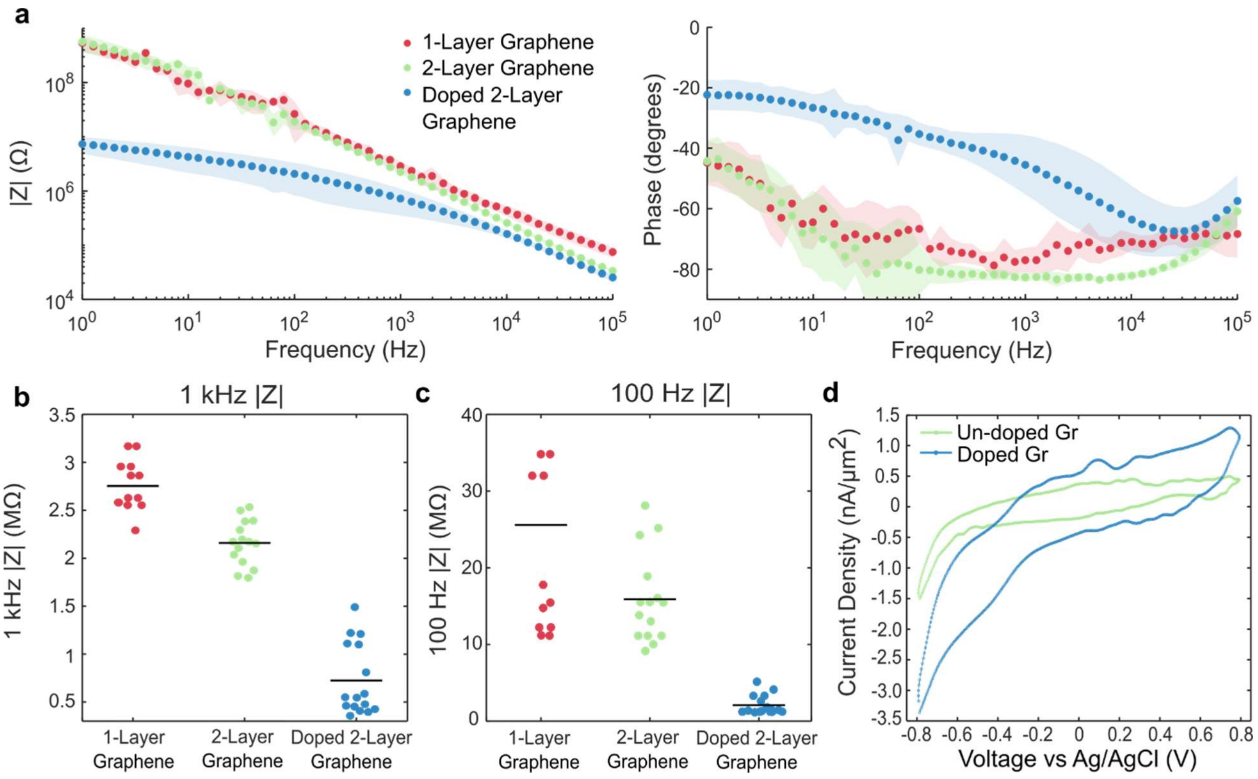


**Supplementary Fig. S2 | Optimization of graphene processing. a**, Electrochemical impedance spectra for 50 μm x 50 μm graphene μECoG electrodes of three preparations: 1-layer graphene, 2-layer graphene, and $HNO_3$-doped 2-layer graphene. Impedance magnitude significantly decreased with doping, particularly at frequencies below 1 kHz. Noise in the low-frequency range for the un-doped graphene samples is attributed to the high impedance modulus. Shaded bounds represent standard deviations. **b**, Impedance magnitude at 1 kHz for the three device preparations. Horizontal black bars indicate means. **c**, Impedance magnitude at 100 Hz for the three device preparations. Horizontal bars indicate means. **d**, Cyclic voltammograms showing superior $CSC_C$ in an $HNO_3$-doped 2-layer graphene electrode, as compared to an un-doped 2-layer graphene electrode ($CSC_c$ is 22.4 μC/cm$^2$ and 15.9 μC/cm$^2$, respectively). The scan rate was 500 mV/s.

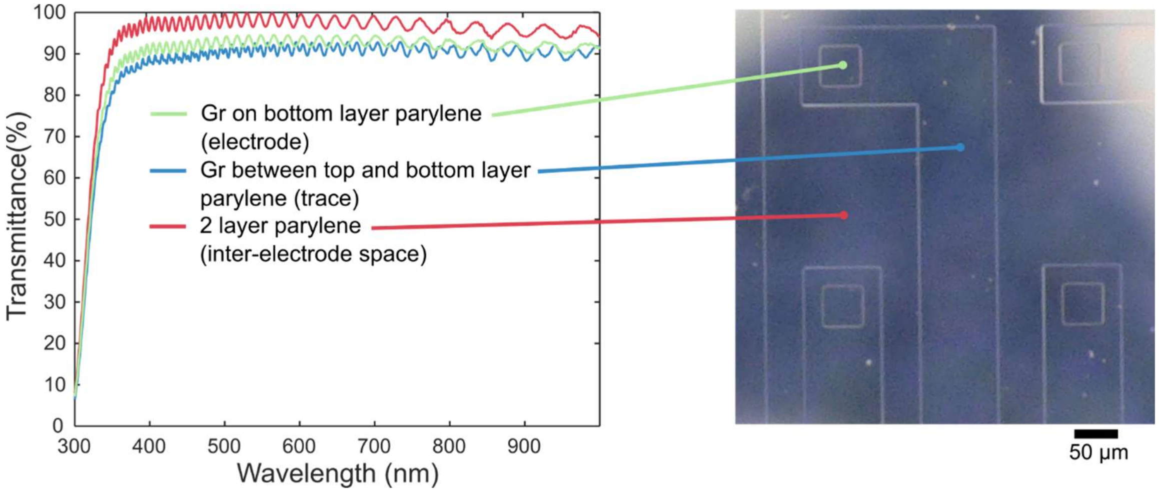


**Supplementary Fig. S3 | UV-Visible spectroscopy.** Optical transmission spectrum for different locations in the graphene μECoG array. Overall, the device is >90% transparent across the visible to near infrared spectrum. The ringing behavior in the spectra is attributed to the parylene-C.

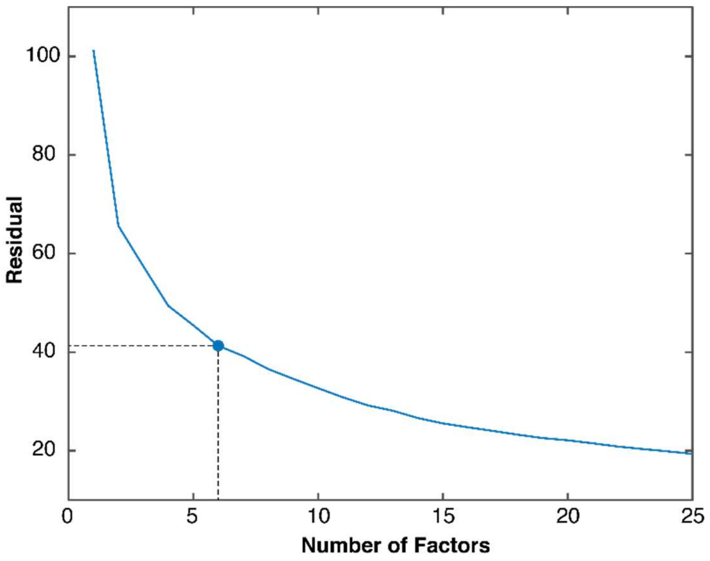


**Supplementary Fig. S4 | Variance explained by number of factors**. Residuals for NMF fits after optimization for preset factor numbers ranging from 1 to 25. The final number of factors (***k*** = 6) was selected as a trade-off between maintaining a low number of factors and achieving a low residual variance.

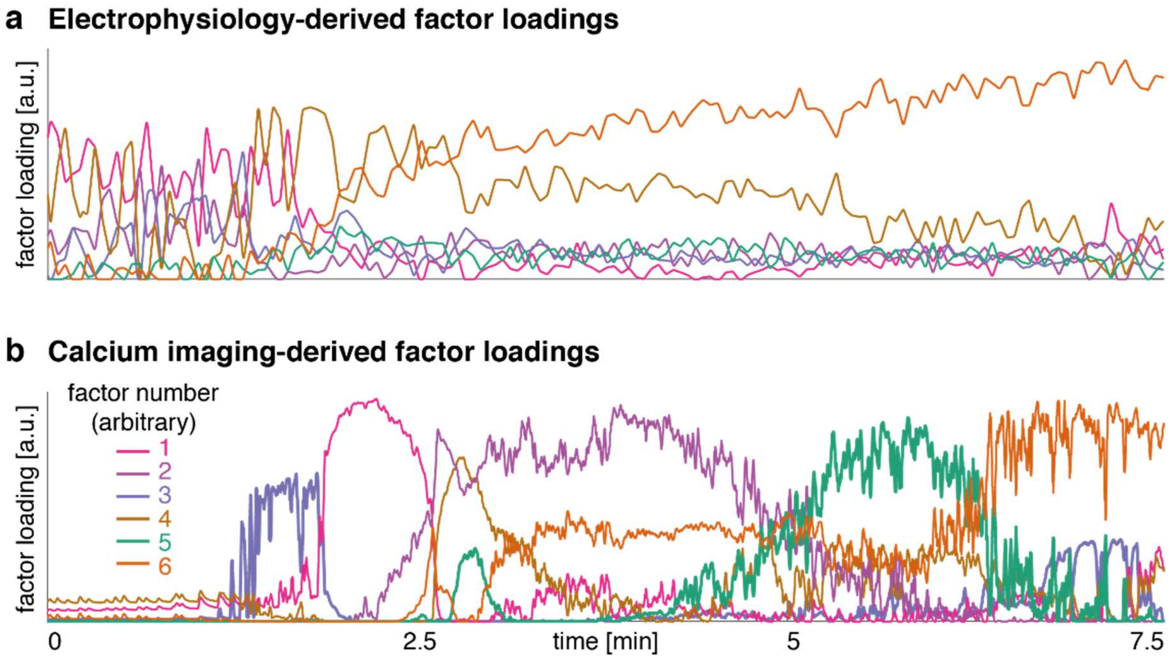


**Supplementary Fig. S5 |** Factor loadings (i.e. ***H***-matrix) derived from non-negative matrix factorization. **a,** Results derived from matrices containing electrophysiology-derived data features only show transition into seizure and some progressive changes, but no clear sequential states. **b,** Factor loadings derived from calcium imaging features only shows sequential states similar to those of the combined analysis presented in the main text.

**Supplementary Video S1 | Multimodal recording during seizure onset.** Onset and spread of the seizure activity analyzed in Fig. 3 – Fig. 6. Here, we show the normalized fluorescence video (right), and the relevant data for one selected electrode (blue): fluorescence intensity beneath the electrode, LFP bandpower, spectrogram, and raw signal.
